# Monkeypox DNA correlates with virus infectivity in clinical samples

**DOI:** 10.1101/2022.08.02.502454

**Authors:** Nir Paran, Yfat Yahalom-Ronen, Ohad Shifman, Shirley Lazar, Ronen Ben-Ami, Michal Yakubovsky, Itzchak Levy, Anat Wieder Feinsod, Sharon Amit, Michal Katzir, Noga Carmi Oren, Ariela Levcovich, Mirit Hershman-Sarafov, Alona Paz, Rebecca Thomas, Hadas Tamir, Lilach Cherry-Mimran, Noam Erez, Sharon Melamed, Moria Barlev-Gross, Shay Karmi, Boaz Politi, Hagit Achdout, Shay Weiss, Haim Levy, Ofir Schuster, Adi Beth-Din, Tomer Israely

## Abstract

In light of the alarming global spread of Monkeypox and the lack of data regarding virus infectivity in clinical specimens, we examined virus infectivity and its correlation with PCR results, the currently available diagnostic tool. Due to the high risk of Monkeypox virus infection, only a limited number of approved BSL-3 laboratories employed with vaccinated staff, including ourselves, are capable of determining Monkeypox virus infectivity. We show strong correlation between viral DNA content and virus infectivity in clinical specimens. Moreover, we define a PCR threshold value which correspond to non-infectious virus, and suggest that our data can be translated into informed decision-making regarding risk assessment, protective measures and guidelines for Monkeypox patients.

---

Monkeypoxvirus (MPXV) is a zoonotic double-stranded DNA virus, of the orthopoxvirus genus within the Poxviridae family [1, 2]. Potential routes of exposure include interaction with wild animals and close contact with sick individuals, as well as contact with infectious fomites (https://www.who.int/news-room/fact-sheets/detail/monkeypox).

MPXV was originally reported in humans in central Africa in 1970, and since then sporadic outbreaks have been reported mostly in central and west Africa. Sporadic MPXV outbreaks in non-African countries have been reported since 2003. In 2018, MPXV of the west-African clade was isolated in both Israel, the United Kingdom and Singapore [3, 4].

In early May 2022, a first confirmed case of MPXV was reported in the United Kingdom. Since then, increasing numbers of cases, attributed to the MPXV West African clade, have been reported in non-endemic countries, currently amounting to over 22,485 cases worldwide (As of July 29^th^, 2022, https://www.cdc.gov/poxvirus/monkeypox/response/2022). As of now, disease is mostly affecting men-who-have-sex-with-men (MSM), and the disease is manifested mainly with skin lesions, at multiple sites including genital, oropharyngeal or perianal areas [5].

Currently, the rapid detection and identification of MPXV is based on positive polymerase chain reaction (PCR) results. However, the presence of viral DNA in a clinical sample does not imply on its infectivity. This uncertainty restricts risk assessment regarding potential transmission from confirmed cases, as well as decision making and setting criteria for quarantine and infection control guidelines. Here, we tested the correlation between DNA copies as measured by PCR, and infectious virus as measured by plaque assay, in 43 PCR-positive, MPXV clinical specimens (CT value<40) obtained from 32 patients, including 21 oropharyngeal swabs, 20 lesion exudate swabs, and 2 rectal swabs (Table S1).

By plotting CT values against log plaque forming units (pfu) titers of all samples irrespective of swabs’ type, we show significant negative correlation (Fig. 1a, r=-0.9349, p<0.0001), where low CT values correlates with high viral titers. Thus, high CT values indicate lower infectivity. Of note, due to inherent nature of monkeypox disease that manifests mostly with dermal lesions and less with oropharyngeal lesions, most lesion swabs exhibit low CT values and high viral load, whereas most oropharyngeal specimens represent the opposite.

**Figure 1:**
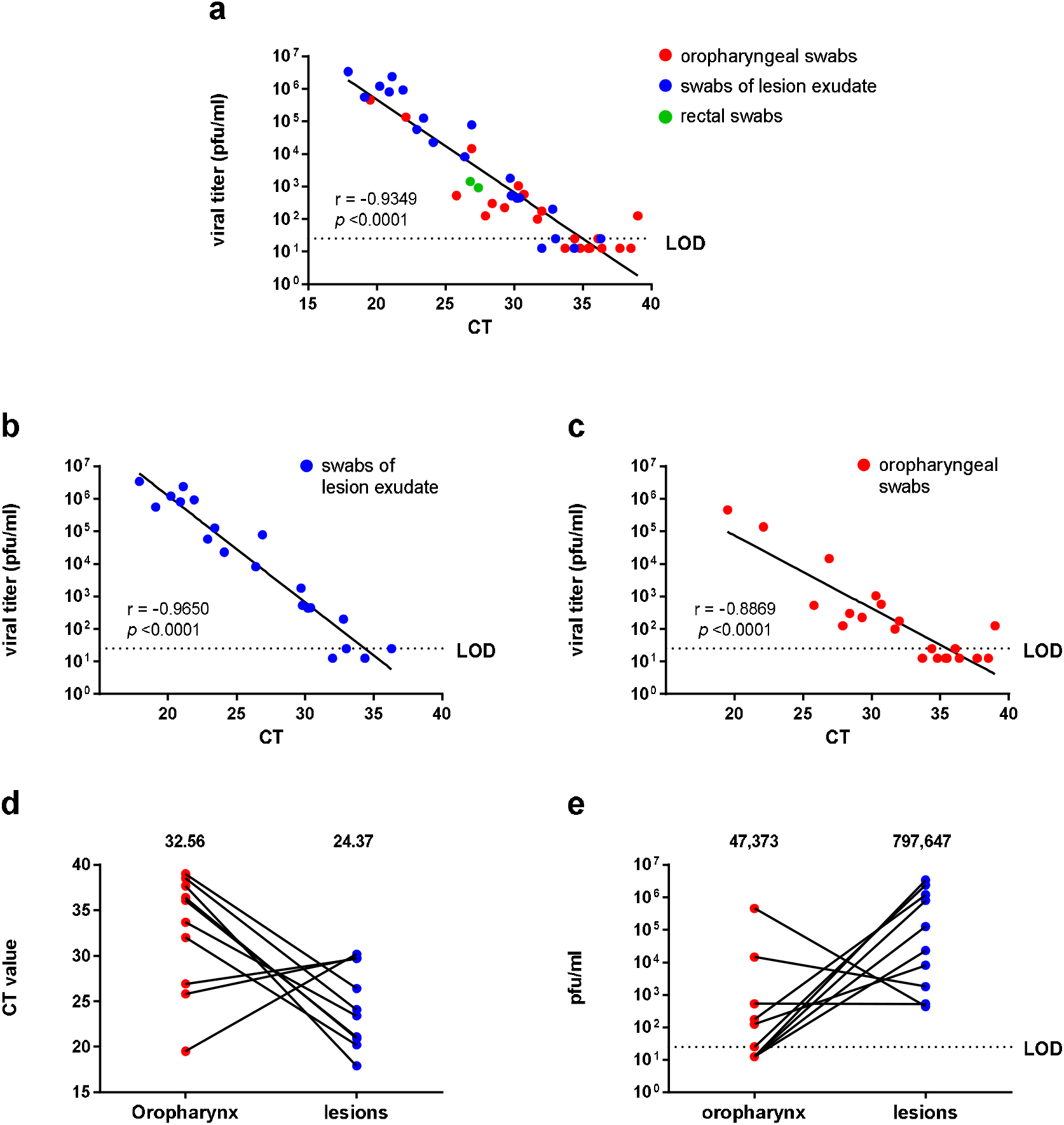
Correlation between CT values and pfu in MPXV clinical specimens: Forty three clinical specimens obtained from 32 patients including 21 oropharyngeal swabs, 20 dermal lesion exudate swabs and 2 rectal swabs were tested for MPXV DNA by real-time PCR and for infectious viral load by plaque assay. (a-c) PCR results represented by cycle threshold (CT) values, and infectious virus load as determined by plaque forming units/ml are plotted. (a) Correlation between CT values and viral load (pfu/ml) in all tested samples, (b) in lesion exudates, and (c) in oropharyngeal swabs. Paired viral DNA (CT values) and infectious virus (pfu/ml) from 10 individuals were tested by (d) PCR and (e) plaque assay. LOD – limit of detection = 25 pfu/ml. Samples below LOD were assigned a value of half LOD (12.5). (a-c) Pearson correlation coefficient was determined for each two variables. (d) Mean CT values and (e) mean viral titers are presented.

In order to examine whether PCR values of oropharyngeal swabs correlate with infectivity similarly to lesion exudate swabs, we correlated CT values and log pfu values for each specimen source separately. We show a significant correlation for both dermal lesions (r = - 0.9650) and oropharynx swabs (r = - 0.8869) (Fig. 1b, c). Thus, CT values strongly predict infectivity for both specimen types. CT values for 9 of the tested oropharynx swabs were relatively high (>33, Fig. 1c). For most of them (6/9) infectious particles were below the limit of detection (LOD=25 pfu/ml), 2/9 were at the LOD, and only 1/9 showed low viral load. This suggests that in oropharyngeal swabs, CT values ≥33 (n=3) often predict no or very low infectivity. Similarly, dermal lesions (Fig. 1b) with CT values greater than 33 were at or below limit of detection, implying a CT value of ≥33 representing a poorly or non-infectious specimen. However, further examination of larger amount of lesion swabs at this range should be examined.

Twenty out of the 43 specimens were paired swabs of oropharynx-lesion from the same patient, thus enabling examination of the CT (Fig. 1d) or pfu (Fig. 1e) pairing per patient for these 2 specimen types. We show that most tested patients present higher viral loads (and lower CT values) in dermal lesions than in oropharynx specimens, suggesting a high infectivity risk from close contact with dermal lesions.

There are several limitations for this study. First, due to the nature of the disease, tested oropharyngeal swabs usually display relatively high CT values, thus conclusions as to lower CT values are lacking. Also, data on virus infectivity from other, poorly available types of clinical specimens such as semen, scabs, and rectal swabs still require further evaluation. Although variation in specimen collection might affect viral or DNA recovery, we suggest that the observed strong correlation (fig.1) despite incorporating specimens from 6 medical centers indicate that this factor is of minor effect. Additionally, both laboratory results and disease progression should be considered for each patient.

Taken together, this work highlights the strong correlation of diagnostic MPXV CT values to virus infectivity, and define a CT threshold (≥33) that predicts poorly or non-infectious specimen. This data may provide valuable and critical criteria for decision making regarding protective measures and guidelines for monkeypox patients and their close contacts. Our data suggest that specimens with CT values greater than 33 should be regarded as having low risk of infectivity.

## Supplementary Material

### Methods

#### Clinical specimens

Forty-three clinical specimens of oropharyngeal swabs (21/43), lesion exudate swabs (20/43), or rectal swabs (2/43) from male patients were tested. Samples were obtained from 32 patients. Ten out of the 32 patients have paired oropharyngeal-lesion exudate swabs. One patient has both oropharyngeal and rectal swabs (Table S1). Swabs were stored in viral transfer media (VTM) containing tubes up to 48h at 4^0^C. Tubes were vortexed for 1 minute before sample collection to either DNA extraction or plaque assay.

#### DNA extraction

DNA was extracted from 0.1 ml of the samples using the QIAamp DNA Mini Kit (Qiagen) using a protocol for blood and body fluids in a QIAcube robot and was eluted in 100 μl H_2_O.

#### MPXV real time PCR

Multiplex real-time PCR assays were adopted from Li et al [Supplementary ref 1] and performed in a 50 μl reaction volume using the SensiFAST™ Probe Lo-ROX kit (BIOLINE). The mix contained viral specific primers (30 pmol per reaction each) and probes (15 pmol per reaction each) detailed below: MPXV generic assay (GE) forward primer (5’-GGAAAATGTAAAGACAACGAATACAG) reverse primer (5 ‘-GCTATCACATAATCTGGAAGCGTA) probe (5’Joe-AAGCCGTAATCTATGTTGTCTATCGTGTCC-3 ‘BHQ1) MPXV West African specific assay (WA) forward primer (5’-CACACCGTCTCTTCCACAGA) reverse primer (5’-GATACAGGTTAATTTCCACATCG) probe (5’FAM-AACCCGTCGTAACCAGCAATACATTT-3 ‘BHQ1)

The PCR was carried out on a QuantStudio 5 real-time PCR system (Applied Biosystems), under the following conditions: 20 sec at 95°C followed by 40 cycles at 1 sec 95°C and 20 sec 60°C [1].

#### Cells and Plaque assay

BSC-1 (CCL-26, ATCC) cells were maintained in DMEM medium containing 10% Foetal Bovine Serum (FBS), MEM nonessential amino acids (NEAA), 2 mM L-glutamine, 100 Units/ml penicillin, 0.1 mg/ml streptomycin, 12.5 Units/ml nystatin (P/S/N), all from Biological Industries, Israel. For plaque assay, tenfold serial dilutions (−1 to −6) from each specimen were performed in MEM medium containing 2% FBS, MEM nonessential amino acids (NEAA), 2 mM L-glutamine, 100 Units/ml penicillin, 0.1 mg/ml streptomycin, 12.5 Units/ml nystatin, all from Biological Industries, Israel. Monolayers of BSC-1 cells in 12-well plates were infected in duplicates, incubated for 1 h at 37°C with 5% CO_2_ incubator. Then 2ml of methylcellulose overlay (MEM containing 0.5% methylcellulose, 2% FBS, 2 mM L-glutamine, 100 Units/ml penicillin, 0.1 mg/ml streptomycin, 12.5 Units/ml nystatin) were added to each well and cells were incubated for 72 h at 37°C with 5% CO_2_ incubator. Plates were fixed and stained with crystal violet, plaques were counted and plaque forming units/ml (pfu/ml) was determined (Fig. S1). Handling and working with MPXV samples were conducted in a BSL3 facility in accordance with the biosafety guidelines of the Israel Institute for Biological Research (IIBR).

#### Statistics

Statistical analyses were performed using GraphPad Prism 6.0. Linear regression was performed on log pfu/ml and CT values. To measure the Correlation between CT and pfu/ml values, we determined the Pearson correlation coefficient (r) for all samples as well as separately for dermal exudate and oropharyngeal swabs. Significance is presented as two-tailed p-value. Mean values are presented for CT and pfu/ml.

**Table S1:** Viral DNA (CT values) and load of infectious virus (pfu/ml) in clinical samples. Forty-three swabs were obtained from 32 male patients. Ten patients have both oropharyngeal and lesion swabs. One patient has both oropharyngeal and rectal swabs.

| patient # | oropharyngeal swabs |  | swabs from lesion exudate |  | rectal swabs |  |
| --- | --- | --- | --- | --- | --- | --- |
|  | CT value | pfu/ml | CT value | pfu/ml | CT value | pfu/ml |
| 1 | 38.5 | 12.5 | 24.1 | 22875 |  |  |
| 2 | 39 | 125 | 26.4 | 8191.7 |  |  |
| 3 | 36.4 | 12.5 | 20.9 | 807500 |  |  |
| 4 | 37.7 | 12.5 | 17.9 | 3400000 |  |  |
| 5 | 26.9 | 14483.3 | 29.7 | 1775 |  |  |
| 6 | 36.1 | 25 | 21.1 | 2400000 |  |  |
| 7 | 33.7 | 12.5 | 23.4 | 126000 |  |  |
| 8 | 25.8 | 535 | 29.8 | 525 |  |  |
| 9 | 32 | 175 | 20.2 | 1209167 |  |  |
| 10 | 19.5 | 458333.3 | 30.2 | 437.5 |  |  |
| 11 | 35.4 | 12.5 |  |  |  |  |
| 12 | 35.5 | 12.5 |  |  |  |  |
| 13 | 34.4 | 25 |  |  |  |  |
| 14 | 34.8 | 12.5 |  |  |  |  |
| 15 | 31.7 | 100 |  |  |  |  |
| 16 | 30.7 | 575 |  |  |  |  |
| 17 | 30.3 | 1050 |  |  |  |  |
| 18 | 29.3 | 225 |  |  | 27.4 | 925 |
| 19 | 28.4 | 300 |  |  |  |  |
| 20 | 27.9 | 125 |  |  |  |  |
| 21 | 22.1 | 136416.7 |  |  |  |  |
| 22 |  |  | 36.3 | 25 |  |  |
| 23 |  |  | 34.4 | 12.5 |  |  |
| 24 |  |  | 33.0 | 25 |  |  |
| 25 |  |  | 32.8 | 200 |  |  |
| 26 |  |  | 32.0 | 12.5 |  |  |
| 27 |  |  | 30.4 | 450 |  |  |
| 28 |  |  | 26.9 | 77500 |  |  |
| 29 |  |  | 22.9 | 57500 |  |  |
| 30 |  |  | 21.9 | 932500 |  |  |
| 31 |  |  | 19.1 | 560000 |  |  |
| 32 |  |  |  |  | 26.8 | 1425 |

**Fig. S1:**
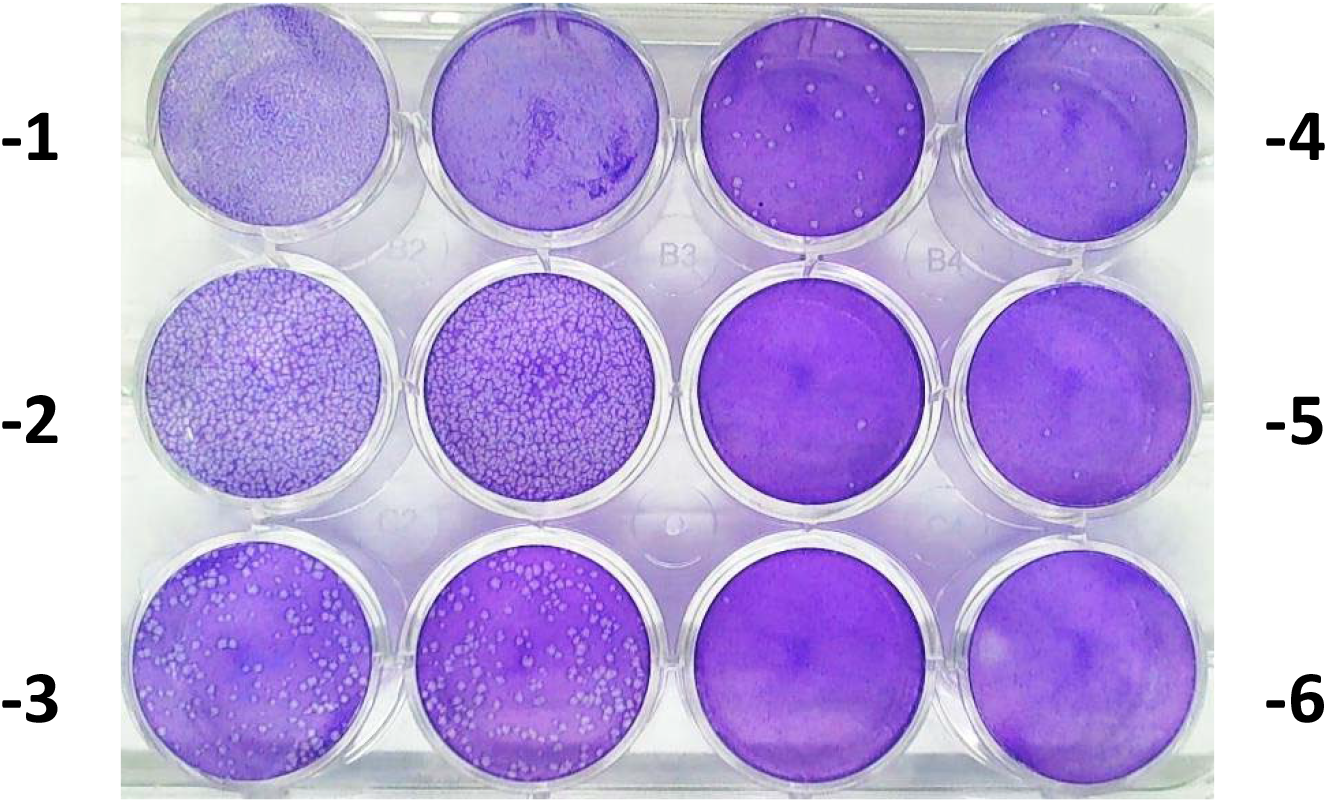
Plaque forming unit (pfu) assay for MPXV clinical samples. Tenfold serial dilutions (−1 to −6) from each specimen were performed. Monolayers of BSC-1 cells in 12-well plates were infected in duplicates, incubated for 1 h at 37°C with 5% CO_2_ incubator. Then 2ml of methylcellulose overlay were added to each well and cells were incubated for 72 h at 37°C with 5% CO_2_ incubator. Plates were fixed and stained with crystal violet. Representative plate is shown.

## References

1. Kmiec, D. and F. Kirchhoff, Monkeypox: A New Threat? Int J Mol Sci, 2022. 23(14).

2. McCollum, A.M. and I.K. Damon, Human monkeypox. Clin Infect Dis, 2014. 58(2): p. 260–7.

3. Erez, N., et al., Diagnosis of Imported Monkeypox, Israel, 2018. Emerg Infect Dis, 2019. 25(5): p. 980–983.

4. Mauldin, M.R., et al., Exportation of Monkeypox Virus From the African Continent. J Infect Dis, 2022. 225(8): p. 1367–1376.

5. Lancet, T., Monkeypox: a global wake-up call. Lancet, 2022. 400(10349):337.

## Supplementary References

1. Li Y, Zhao H, Wilkins K, Hughes C, Damon IK. Real-time PCR assays for the specific detection of monkeypox virus West African and Congo Basin strain DNA. J Virol Methods. 2010 Oct;169(1):223–7. doi:10.1016/j.jviromet.2010.07.012. Epub 2010 Jul 17. PMI: 20643162.

